# Sequence Determines the Switch in the Fibril Forming Regions in the Low Complexity FUS Protein and Its Variants

**DOI:** 10.1101/2021.07.08.451535

**Authors:** Abhinaw Kumar, Debayan Chakraborty, Mauro Lorenzo Mugnai, John E. Straub, D. Thirumalai

## Abstract

Residues spanning distinct regions of the low-complexity domain of the RNA-binding protein, Fused in Sarcoma (FUS-LC), form fibril structures with different core morphologies. NMR experiments show that the 214 residue FUS-LC forms a fibril with an S-bend (core-1, residues 39-95), while the rest of the protein is disordered. In contrast, the fibrils of the C-terminal variant (FUS-LC-C; residues 111-214) has a U-bend topology (core-2, residues 112-150). Absence of the U-bend in FUS-LC implies that the two fibril cores do not coexist. Computer simulations show that these perplexing findings could be understood in terms of the population of sparsely-populated fibril-like excited states in the monomer. The propensity to form core-1 is higher compared to core-2. We predict that core-2 forms only in truncated variants that do not contain the core-1 sequence. At the monomer level, sequence-dependent enthalpic effects determine the relative stabilities of the core-1 and core-2 topologies.

**TOC graphic:** 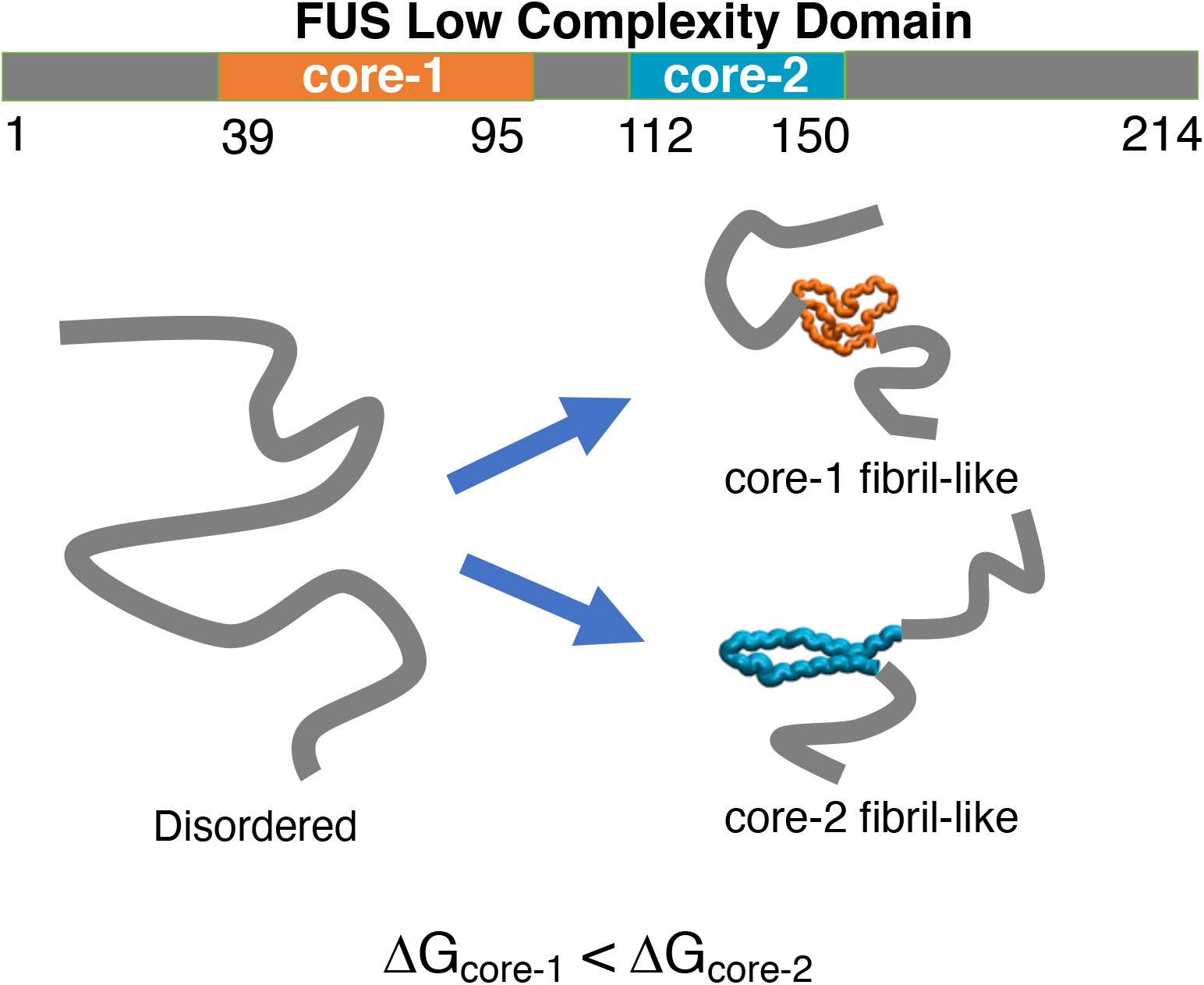

---

Fused in sarcoma (FUS), a RNA-binding protein, has garnered considerable attention in recent years, not only because it is implicated in neurological disorders and biological functions, ^1–3^ but also because it serves as a prototype for elucidating the principles underlying condensate formation in low-complexity protein sequences. ^4–6^ The N-terminal region of the 526-residue FUS protein houses the low-complexity (LC) domain, rich in QGSY repeats. The FUS-LC domain (residues 1-214) is intrinsically disordered, and the ground state likely behaves as a random-coil. On the other hand, the C-terminal end of the full-length FUS is ordered and is involved in protein as well as RNA binding.^7^

Due to its relevance in liquid-liquid phase separation (LLPS) and fibril formation, the FUS-LC domain has been the subject of several experimental biophysical studies, ^4,8–21^ and simulations.^22–30^ The experimental findings that are relevant to our study, may be summarized as follows: (i) A combination of fluorescence microscopy, and solution-state NMR of a truncated version (residues 1–163) of FUS-LC indicate that it is devoid of persistent structure, both in the monomer as well as in the liquid droplet state.^8^ In contrast to a previous report based on X-ray crystallography and negative stain Transmission Electron Microscopy (TEM)^10^ experiments, no signatures of hydrogel formation were immediately apparent in the condensed phase. (ii) Using solid-state NMR, Tycko and coworkers^9^ showed that a 50 *µ*M solution of FUS-LC does form fibrils, characterized by an S-shaped ordered region involving only residues 39-95. In the S-shaped structure (core-1), the monomers are arranged as parallel *β*-strands (Figure 1). The rest of the residues are not sufficiently ordered to be resolvable in experiments. They form a fuzzy coat surrounding the fibril region. Based on the observation that NMR cross-peaks were reproducible in independent measurements, it was concluded that FUS-LC did not show any evidence of fibril polymorphism. (iii) Surprisingly, in a follow-up study, Tycko and coworkers^31^ demonstrated that a truncated version of FUS-LC, devoid of the residues forming the core-1 fibril, and comprising only the C-terminal region of the protein (FUS-LC-C; containing residues 110-214) also forms a fibril, with an *entirely different morphology* (Figure 1). In this structure, residues 112-150 form a U-shaped fibril (core-2), while the rest of the protein is disordered. To rationalize their findings, it was argued^31^ that the formation of core-2 is suppressed in FUS-LC because in such a structure the N-terminal and C-terminal disordered segments, constituting the fuzzy coat, would come in close proximity due to the U-bend topology, resulting in a decrease in the effective available volume. In other words, it was conjectured that the lack of core-2 for FUS-LC could be attributed to an entropic effect.

**Figure 1:**
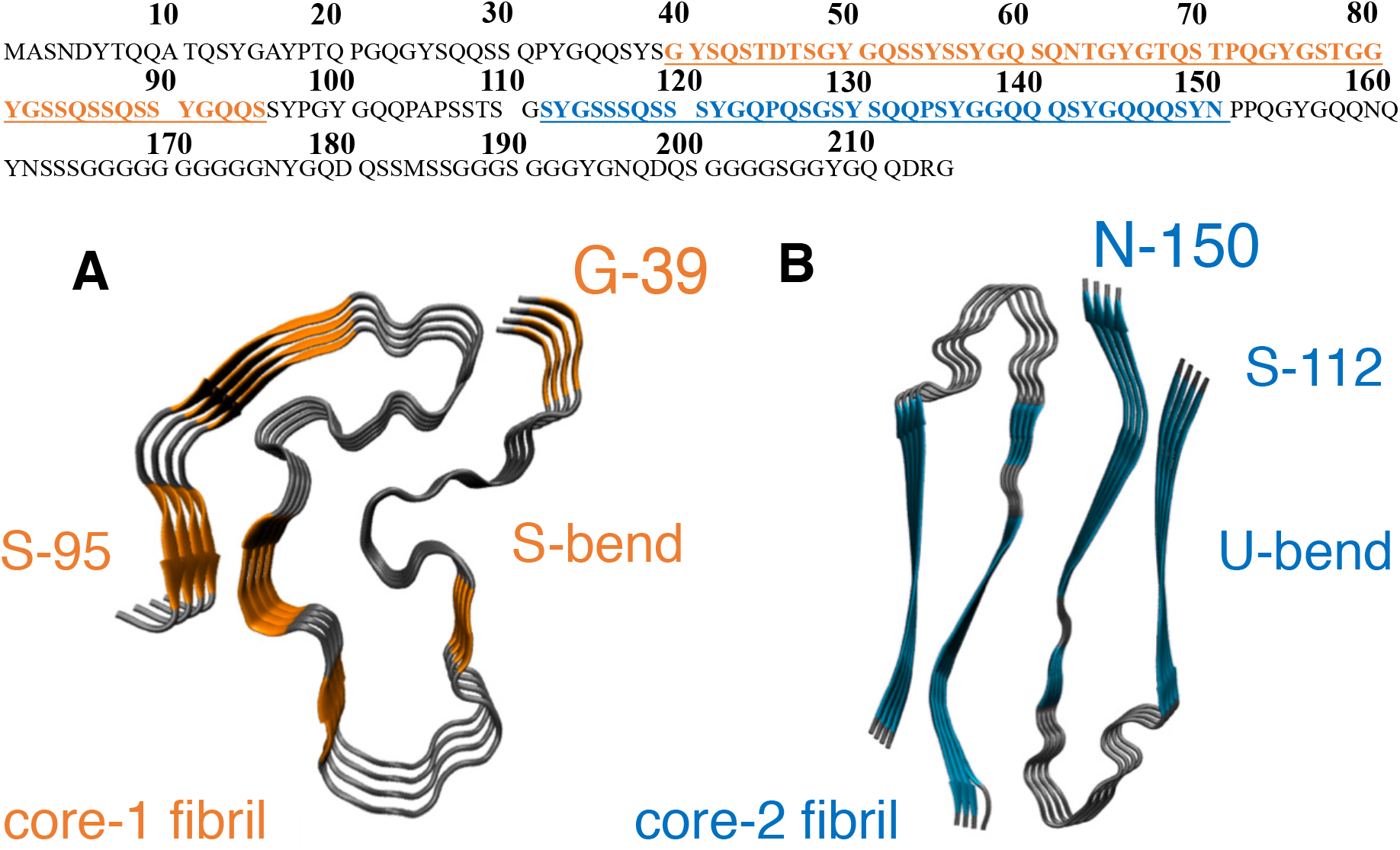
Top panel: The FUS-LC sequence, denoted using a one letter code for the amino-acids. Residues 39-95, which form the core-1 fibril, are highlighted in orange, and those within core-2 (112-150) are highlighted in blue. (A) The experimental structure for the core-1 fibril, exhibiting a S-bend morphology (PDB ID: 5W3N). ^9^(B) The experimental core-2 fibril structure^31^ exhibiting a U-bend morphology (PDB ID: 6XFM).

Further impetus into this debate, which centers around the probable structures (liquid-droplet, hydrogel, or fibril) of FUS-LC, and how transitions between the different forms could occur, comes from the recent work based on magic-angle spinning NMR spectroscopy and imaging.^11^ This study suggests that the liquid-droplet state may represent a metastable state of FUS-LC, and gradual aging could lead to the fibril structure, the likely thermodynamic end-product.

In light of the experimental results summarized above, the following questions arise: (a) Why does core-2 not form in FUS-LC? The non-existence of core-2 is indeed puzzling because the solubility of an N-terminal construct derived from the FUS-LC sequence (FUS-LC-N2; residues 2-108), which forms core-1 fibrils, was shown to be similar to FUS-LC-C. Although the rationale^31^ based on purely entropic arguments is likely to be correct on the scale of fibrils, there could be subtle enthalpic effects that manifest more readily at the monomer level. Somewhat indirect evidence for such a possibility comes from a previous study, which observed that phosphorylation of six residues within core-1 strongly affected the recruitment of solvated polymers in hydrogel droplets. In contrast, phosphorylation of peripheral residues had no effect.^9^ (b) Does a truncated variant of FUS-LC-N (residues 1-163) have the propensity to form the fibril structure? Here, we provide some plausible answers to these questions using computer simulations based on an accurate sequence-specific coarse-grained model for intrinsically disordered proteins.^32,33^ We probe the conformational landscape of the FUS-LC monomer, and its variants (the truncated and the phosphorylated forms) to ascertain if the free energy excitations within the monomer conformational ensemble (MCE) could provide insights into the conundrum that prevails regarding the feasible structures and polymorphism of FUS-LC.

Our work is based on the premise that the free energy spectrum of the monomer, and the emergent dynamical heterogeneity of the MCE, have robust links to the early events in the assembly cascade, leading to the formation of a liquid-droplet, hydrogel, and ultimately the fibril state. ^34,35^ The harbingers of self-assembly (referred to as N^∗^ states) present within the MCE are sparsely populated (usually ≈ 2% or less), and are challenging to characterize even with advanced NMR techniques.^36^ The N^∗^ states usually have some elements of structural order (akin to fibrils) interspersed with fully disordered segments, which predisposes them to coalesce with other assembly-competent conformations to form oligomers of different sizes. In previous studies, we have shown that the N^∗^ concept accounts for the sequence-specific fibril formation time scales and provides a microscopic basis for fibril polymorphism. ^33,37,38^ Within the MCE of FUS-LC, only a very few N^∗^ conformations having core-2 fibril-like features are found. On the other hand, N^∗^ states corresponding to core-1 appear with higher probability despite having a more complex morphology than core-2. Our observation suggests that, in addition to entropic effects, sequence-specific enthalpic contributions, apparent at the monomer level, contribute to the free energy difference between the fibril morphologies. The relatively low population of N^∗^ state within the MCE contributes to the overall destabilization of the core-2 polymorph.

## Fingerprints of fibrillar order within the MCE

The equilibrium free energy landscape of the FUS-LC monomer is largely featureless like other IDPs, and lacks discernible metastable states (Figure S2). To quantify the similarity with respect to the fibril-state, we compute the structural overlap parameter,^39^ *χ* (see the Supplementary Information for definition), between a conformation sampled along the simulation trajectory and a monomer unit within the experimental fibril structure. ^9^

The distributions of the overlap parameter, P(*χ*)s, with respect to the core-1 and core-2 fibril structures are shown in Figure 2. In both cases, the *χ* distributions peak at relatively low values, suggesting that the MCE is conformationally heterogeneous, and on average, bears little or no resemblance to the fibril states. However, a careful structural analysis of the sub-ensembles corresponding to the tails of the *χ* distributions does reveal the existence of a sparse population of fibril-like (N^∗^) conformations.

**Figure 2:**
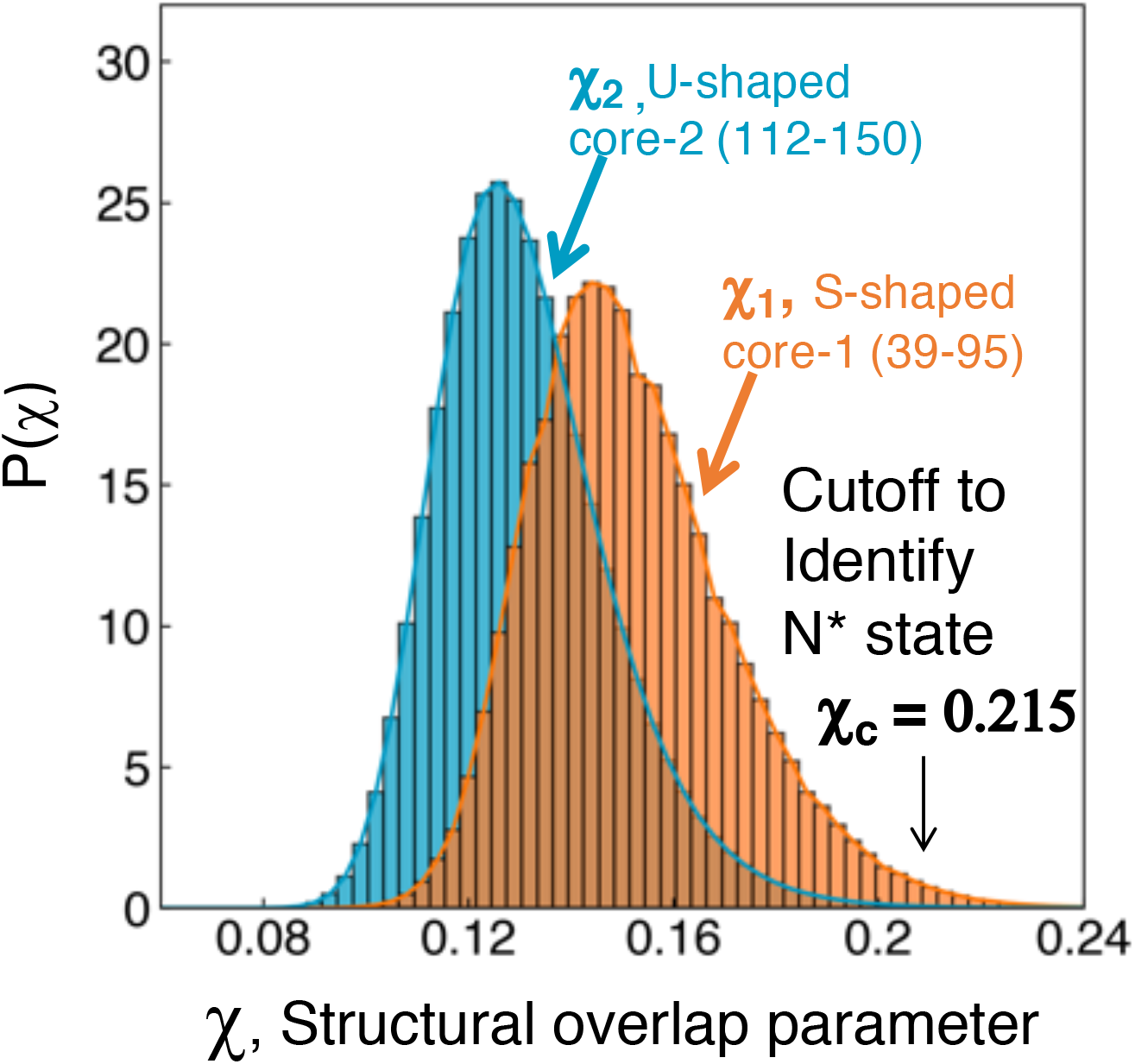
The distribution of the structural overlap parameter, *χ*, for the FUS-LC sequence; *χ* (defined in Eq. S9 in the SI) was computed with respect to a monomer unit within the experimental fibril structures. For the core-1 fibril, the solid-state NMR structure (PDB ID: 5W3N) for the S-bend morphology^9^ is used as the reference. For the core-2 fibril, the cryo-EM structure (PDB ID: 6XFM) the reference structure is the U-bend morphology.

We define N^∗^ states as the ensemble of conformations for which *χ* ≥ *χ*_*c*_, where *χ*_*c*_ separates the basin of attraction of fibril-like structures from the disordered states. The relative positions of the *χ* distributions (Figure 2) suggest that the population of core-1 type N^∗^ conformations is higher than those with core-2 type structures, for any reasonable choice of *χ*_*c*_. This result is unexpected because from a purely entropic consideration we would expect that the formation of the complex, longer S-bend shape is less likely than the simpler, shorter U-bend morphology (see the SI for further discussion of the relative stabilities of core-1 and core-2). But our simulations show the opposite trend, suggesting that sequence-specific effects must play a role.

The residue-residue contact maps (Figures 3(a) and (c)), reveal that the N^∗^ states (defined for *χ ≥ χ*_*c*_ = 0.215) share a few structural features in common with the fibril-state. In particular, the S-bend and U-bend morphologies of the two cores are captured, showing that the choice of cutoff is reasonable. Nonetheless, the imperfect structural alignment with respect to a monomer unit from the fibril also implies that the N^∗^ states retain some degree of disorder. This is not surprising because the stability of a monomer in the fibril arises from favorable interactions with its neighbors. Furthermore, N^∗^ states need to be fluid enough to recruit additional monomers and form oligomers of different sizes. We conclude by remarking that the simulations were performed in the absence of potentials biasing the sample towards the N^∗^ state. In this sense, the emergence of fibril-like conformations is a genuine outcome of the model.

**Figure 3:**
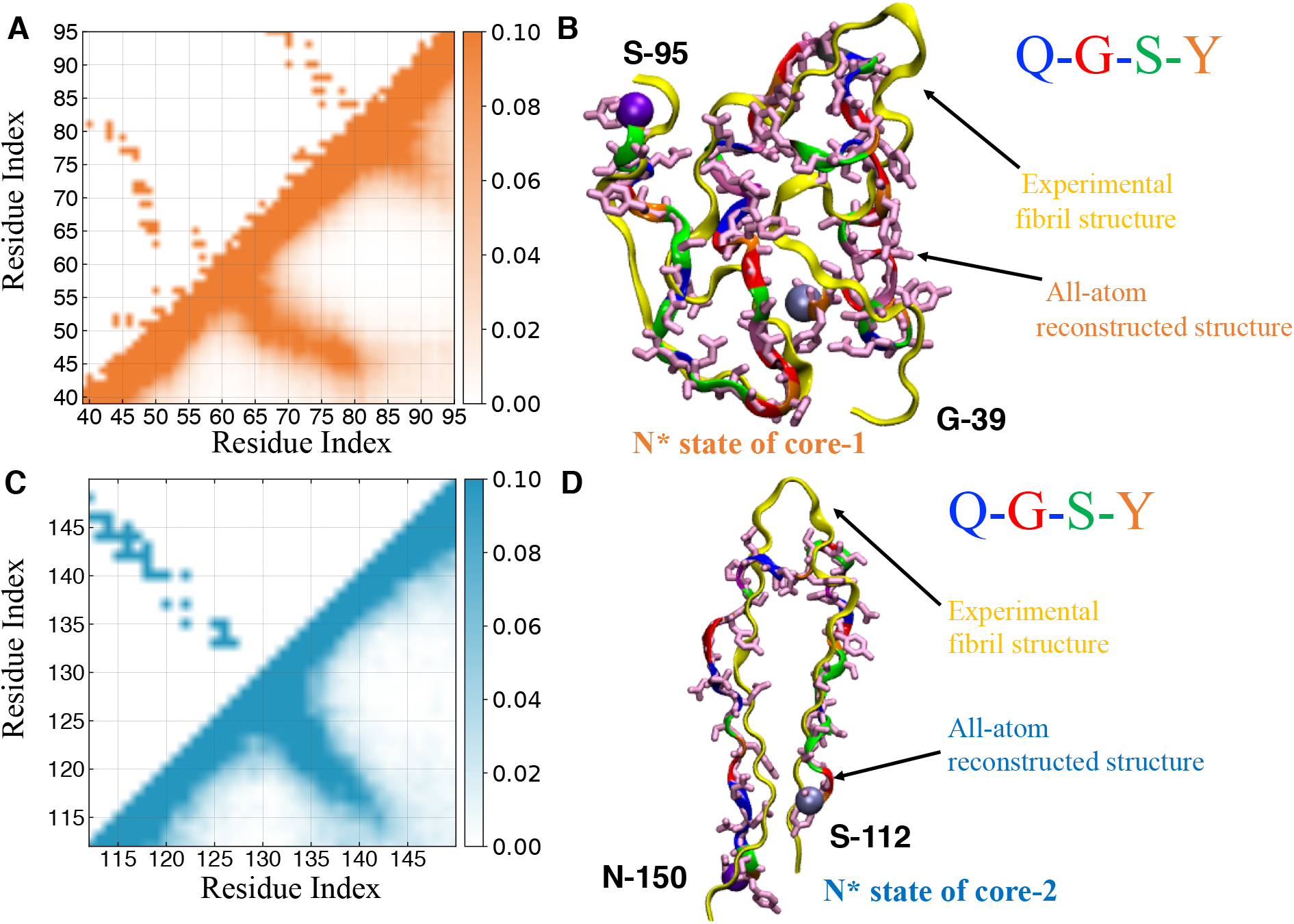
(A) and (C) Contact maps for the N^∗^ ensemble identified from simulations (lower triangle), and a monomer unit from the experimental fibril structure (upper triangle). A structure from the N^∗^ ensemble is shown superimposed on a monomer experimental fibril structure (shown in yellow). For a systematic comparison, all-atom structures were generated from the coarse-grained snapshots (see Supplementary Information for further details). The N^∗^ ensemble corresponding to the core-1 fibril is shown in (B), for core-2 is shown in (D). It is clear that the N^∗^ conformations share considerable similarity with the fibril counterparts.

## Truncation effects

In order to quantify the effects of truncation of sequence on the probabilities of core-1 and core-2 formation we calculated Δ*P*(*χ*_*i*[*α*]_) = *P*(*χ*_*i*[*α*]_) − *P*(*χ*_*i*[*FUS*−*LC*]_) (*i*=1 and 2 are labels for core-1 and core-2, respectively and α is a label for the simulated FUS constructs). It is evident from the difference between the *χ*-distributions of a FUS variant and the WT FUS-LC (blue and green histograms in Fig. 4) that there is practically no effect of the two N-terminal truncations (FUS-LC-N2 (residues: 1-108); FUS-LC-N (residues 1-163)) on the overall conformational heterogeneity. These findings imply that the population of N^∗^ states (and by inference, the propensity to form fibrils) is unaffected by sequence truncations. Our observation is consistent with experimental findings, ^31^ which showed that the FUS-LC-N2 assembles into fibrils similar to those formed by the full-length FUS-LC. Our results also suggest that the fibril state must be the thermodynamic end-state for the FUS-LC-N as well, a prediction that could be validated using solid-state NMR experiments (Figure S5).

**Figure 4:**
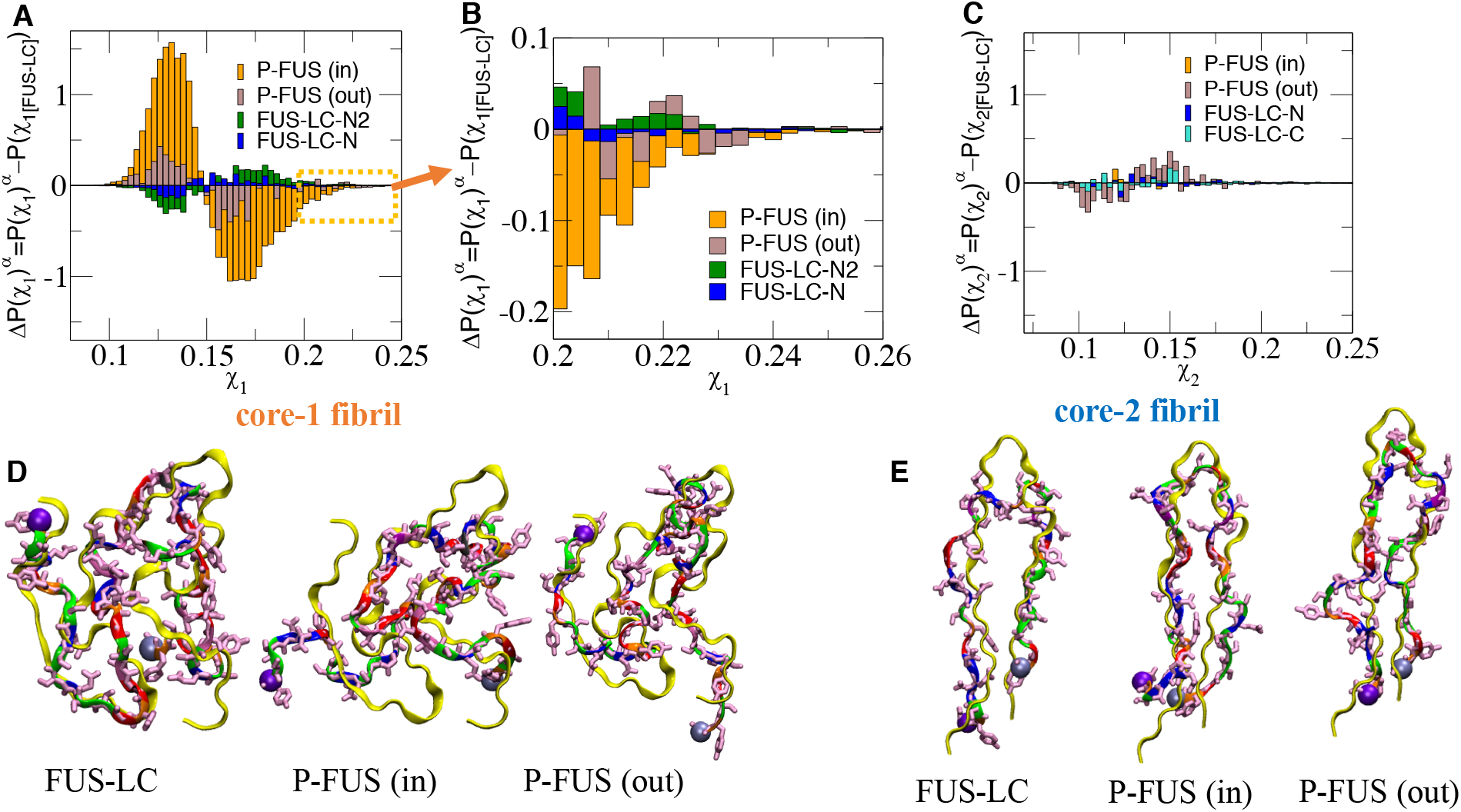
The difference in the probability distribution, P(*χ*), of the structural overlap parameter. (A) Difference plots, Δ*P*(*χ*_1[*α*]_) = *P*(*χ*_1[*α*]_) − *P*(*χ*_1[*FUS*−*LC*]_), for truncated and phosphomimetic mutants where *α* labels the four labeled variants. The four variants are: (i) FUS-LC-N (residue 1-163) (blue); (ii) FUS-LC-N2 (residue 1-108)(green); (iii) P-FUS (in) (orange) corresponds to monomer with phosphorylation inside the core-1 region; (iv) P-FUS (out) (brown) represents the FUS-LC mutant with phosphorylation outside the core-1 region (B) Zoom-in version of figure A showing that the FUS-LC with the mutant with phosphorylation inside the core region (P-FUS (in)) significantly reduces the population of fibril-like state. (C) Same as (A) except Δ*P*(*χ*_2[*α*]_) = *P*(*χ*_2[*α*]_) − *P*(*χ*_2[*FUS*−*LC*]_) for the four labeled variants corresponding to core-2 are plotted. We used a chi-square test to verify that in panels (A) and (C) the only construct showing a *P*(*χ*) significantly different from FUS-LC is P-FUS (in) for core-1 (p-value = 0). This analysis was conducted using the bin-size shown in panels (A) and (C), grouping bins with fewer than 5 data points, and considering only decorrelated conformations (i.e. separated by about twice the correlation time). (D) and (E) The left, central and right panels in fig. (D) and (E) compare the overlap of the N^∗^ state obtained using simulation with the experimental fibril structure (core-1 in (C) and core-2 in (D)) for FUS-LC, FUS-LC with phosphorylation mutant inside the core, and FUS-LC mutant outside the core. The experimental fibril structures are shown in yellow.

## Effects of phosphorylation

Using a combination of solution-state NMR and microscopy, it was shown that both phosphorylation and phosphomimetic variations of FUS-LC tend to disrupt aggregation as well as phase separation. ^40,41^ In particular, phosphorylation within core-1 sites significantly reduced the propensity of FUS-LC monomers to bind to hydrogel droplets, while phosphorylations outside the fibril core had little impact. To simulate a variant of FUS-LC (denoted as P-FUS (in)), where five Serine residues within core-1 (S42, S54, S61, S84, S87) and a Threonine (T68) are phosphorylated, we replaced the phosphorylated residues by Glutamic acid (see SI for details). The difference between the distributions, Δ*P*(*χ*_1,[*α*]_) (brown and orange histograms in Fig. 4), have prominent features. The peak at low values of *χ*, and the dip beyond *χ*_*max*_ at which *P*(*χ*) (see Fig. 2) is a maximum suggest (orange histograms in Fig. 4 (A) and (B)) that the structural similarity of the MCE relative to the fibril state reduces greatly. Phosphorylation also seems to destabilize some of the key interactions within the N^∗^ state. For the P-FUS (in) variant, not even the conformation structurally closest to the core-1 fibril exhibits the complete S-bend morphology (middle structure in Fig. 4D). The features in Δ*P*(*χ*_1,[*α*]_) are less prominent (purple histograms in Fig. 4 (A) and (B)) for the construct P-FUS (out), in which T11, T19, and T30 and S112, S117, and S131 outside core-1 were phosphorylated. This suggests that the effect of phosphorylation is less disruptive in this case compared to P-FUS (in). Most of the conformations within the N^∗^ ensemble seem to retain the overall S-bend topology (Fig. 4 (D)), which accords well with experimental findings. The reduced aggregation propensity of the phosphorylated variants could be attributed to the enhanced free energy gap between the disordered ground state and the N^∗^ state.

In contrast, the distributions Δ*P*(*χ*_2,[*α*]_) for the four variants of FUS-LC displayed in Fig. 4 (C) show that neither truncation of the sequence, nor phosphorylation affect core-2 formation. Interestingly, representative structures from the N^∗^ ensembles of the phosphorylated variants retain a high degree of structural similarity to the core-2 fibril (Fig. 4 (E)), suggesting that sequence perturbations may be better tolerated within the U-bend morphology.

The results in Fig. 4 (C) and (E) show that the formation of core-2 is not precluded by the formation of core-1. The FUS-LC-C construct, which was used to determine the existence of the alternate fibril structure, ^31^ is practically indistinguishable in terms of *p*_*N*_∗ (equilibrium population of the N* state) from the full length FUS-LC. We infer this result from the small difference observed in the *χ* distributions and probability of forming *N*^∗^ states in FUS-LC and FUS-LC-C (Figure 4C). Taken together, the simulations using FUS-LC and a number of variants show that, although co-existence of core-1 and core-2 is not ruled out based on thermodynamic considerations, the probability of formation of core-2 structure in FUS-LC is sufficiently decreased that it cannot be easily detected. In the absence of the core-1 sequence, when there is no competition, core-2 fibrils can form, and its signature is present even at the monomer level.

We ought to stress that the higher propensity for the formation of the core-1 fibril over core-2 in FUS-LC is a remarkable finding. The entropic penalty associated with the formation of the S-bend motif is certainly larger than for the U-bend. The 50-residue core-1 structure could be thought of as a juxtaposition of two 35-residue U-bend cores sharing a common edge. Hence, the geometrical requirement to form core-2 are included, but not sufficient to form the core-1 structure. For an athermal homopolymer, lacking any sequence-specificity, the population of core-2 type conformations would be substantially higher than those compatible with the core-1 topology. Therefore, entropic effects alone cannot account for the relative stabilities of the core-1 and core-2 structures at the monomer level. We tested our assertion using the following calculation (see SI for further details): the free energy difference between core-1 and core-2-type structures within the MCE is, Δ*G*_12_ = *G*_1_ − *G*_2_ ≈ −1.47 kcal/mol. The difference in energy, Δ*U* = *U*_1_ − *U*_2_, ≈ −2.55 kcal/mol; the entropy difference between the two states would be *T*Δ*S* ≈ −1.08 kcal/mol. In other words, the entropy of core-1 would be less than core-2. Hence, subtle enthalpic effects, which are encoded in the sequence of FUS-LC, determine the stabilities of the aggregation-prone states of the monomer and could be important in understanding the microscopic basis of fibril polymorphism or its absence.

The intermolecular interactions, which are not accounted for in our monomer simulations, could contribute to the conformational selection of the stable fibril morphology. However, the empirical argument introduced by some of us,^33,37^ which relates the time scales of fibril formation to the population of aggregation-prone conformations (N*) in the monomer energy landscape (see below), does not depend on the precise details of the intermolecular interactions between protofibril units within a fibril structure. The essence of the N* theory is that for aggregation to occur the signatures of fibril-like states must already be present within the monomer conformational ensemble, regardless of the ultimate fibril morphology.

## Conclusions

Our simulations answer the questions posed in the Introduction: (a) Why does core-2 not form in FUS-LC? Our findings suggest that within the MCE there is a higher population of assembly-competent (N^∗^) states corresponding to the core-1 structure, compared to the core-2 structure. The lower free energy gap, Δ*G*, between the ground and the N^∗^ state, further implies that the initial events in the aggregation cascade are likely to be more conducive to the core-1 formation, rather than core-2. In prior works, ^33,37^ a correlation between the probability of forming N* states and the time required for forming fibrils was demonstrated. Based on this observation, we propose that the higher probability of forming N* states exhibiting characteristics of core-1, as opposed to core-2, implies a kinetic preference for the formation core-1 fibrils over core-2 fibrils. (b) Does the FUS-LC-N (Residues 1-163) form a fibril ? Our simulations show that there is no reason (apart from a kinetic one) why the FUS-LC-N construct should not form a fibril. We propose that this construct should form a fibril having the core-1 morphology, because irrespective of truncation or phosphorylation, the core-1 structure is always thermodynamically favored.

A similar behavior (core-1 formation in one segment that prevents the formation of core-2) is found in the low complexity TAR DNA-binding protein 43 (TDP43). ^42^ A common structural characteristic of FUS-LC and TDP43 is the following. The segments that are ordered involve residues that are consecutive to each other. Core-1 in FUS-LC is comprised of residues 39-95, whereas in TDP43 it stretches from 311-360. More importantly, all the interactions within core-1 in these IDPs are only between residues that are consecutive along the sequence. There are no stable contacts between residues in the core-1 region and those that are upstream or downstream along the sequence. This is in contrast to *α*-synuclein, for example, in which the fibrils contain contacts between residues that are well-separated (long range) from each other along the sequence. In other words, the chain folds upon itself in the fibril. The differences may arise because *α*-synuclein is not a low complexity sequence.

We conclude with a few testable predictions. (i) Our findings for FUS-LC and previous theory^37^ allow us to qualitatively predict the kinetics of fibril formation. The first passage times for fibril formation starting from the monomer are related to the population of N^∗^ states as,^37,43^ *τ*_*fib*_ *∝* exp(−*Cp*_*N*_∗), where the population, *p*_*N*_∗, of N^∗^ states is expressed as a percentage, and the value of *C* ≈ 1. The empirical relationship given does not account for the multiple pathways that may be involved in the aggregation cascade. Nonetheless, it has been exploited in previous studies to predict the experimental trends in fibril formation times nearly quantitatively.^33,38^ Based on the relative populations of the N^∗^ states within the MCE of FUS-LC, we surmise that the core-1 fibril would form faster than the core-2 fibril. A higher value of *p*_*N*_∗ for core-1 also implies a smaller free energy gap, Δ*G*, between the ground and the assembly-competent state. Our prediction could be (at least qualitatively) tested by obtaining the rates of fibril formation in various FUS-LC constructs.

(ii) At a first glance, our observations leave us with two complementary explanations for the dominance of core-1 over core-2 in FUS-LC fibrils. The argument based solely on the entropy of the disordered tails, which would hold provided stable oligomers (perhaps even a dimer) first forms, does explain the absence of core-2 structure in the FUS-LC fibril. Our study suggests that the enhanced stability of core-1 fibril manifests itself already at the monomer level, and it is predominantly a consequence of sequence-specific energetic effects. Thus, on small sizes (monomers and oligomers), stability is determined by enthalpy whereas on the length scale of the fibril, entropy is likely to be more dominant. In order to test the relative importance of entropy and enthalpy it would be crucial to investigate whether a FUS-LC-N (Residues 1-163) construct forms a fibril, and if so determine its structure. We predict that for FUS-LC-N, core-1 fibrils will form exclusively, and that there should be no difference relative to the full-length FUS-LC construct. The destabilization of core-2 fibrils due to the disordered tails is likely to be weaker in the FUS-LC-N construct, because in this construct the C-terminal tail is shortened by about 50 residues. From the perspective of core-2, FUS-LC-N should be similar to FUS-LC-C: only one unstructured tail (N-terminal for FUS-LC-N, C-terminal for FUS-LC-C) stems from the core of the fibril.

(iii) It would be interesting to ascertain if in a solution of FUS-LC seeded with FUS-LC-C fibrils, core-2 structures could emerge under appropriate conditions. We anticipate that such a setup would significantly reduce the nucleation barrier for core-2 formation. However, if the disordered tails completely prevent the formation of a core-2 fibril for FUS-LC, we would expect seeding to have no impact on fibril growth. This prediction is also testable.

## Acknowledgement

We acknowledge the Texas Advanced Computing Center (TACC) for providing the necessary computing resources. This work was supported by a grants from the National Institutes of Health (GM-107703), National Science Foundation (CHE 19-00033) and a grant from the Welch Foundation (F-0019) administered through the Collie-Welch Regents Chair.

## Supporting Information Available

The details of the model and simulations as well as data analyses are available in the Supplementary Information. This material is available free of charge via the Internet at http://pubs.acs.org/.

## Supplementary Information for

### Computational Methodology

#### SOP-IDP Energy Function

The wild-type FUS LC monomer, truncated and phosphorylated FUS sequences were modeled using the Self-Organized Polymer (SOP) model for intrinsically disordered proteins (IDPs).^S1,S2^ The SOP-IDP has been validated by showing that it quantitatively reproduces the scattering profiles for a diverse range of IDPs,^S1^ as well as the sequence-dependent aggregation behavior of A*β* peptides. ^S2^ In the SOP-IDP model, each amino-acid residue is represented by two interaction sites: a backbone bead (BB) centered on the C_*α*_ atom, and a side-chain bead (SC) centered on the center-of-mass of the side-chain. The coarse-grained interaction potential is:

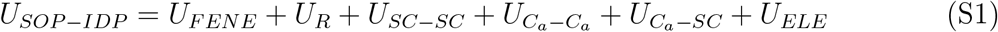

The FENE potential in Eq. S1 accounts for the chain connectivity.

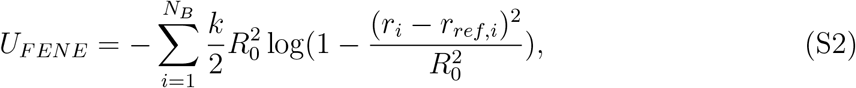

where *r*_*ref,i*_ is the equilibrium distance between the bonded moieties. For a covalent bond between two backbone (*C*_*α*_) atoms, the value of *r*_*ref,i*_ = 2*σ*_*BB*_ where *σ*_*BB*_ is the VDW radius (=1.9 Å) of the *C*_*α*_ atom. If the covalent bond is between a backbone and a sidechain (SC) bead then 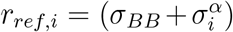. Here, *i* is the residue number and *α* is the type of that residue number. The values of 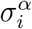 are tabulated in Table S1. In equation S2, *k* is the stiffness of the bond and *R*_0_ is the tolerance for the anharmonic fluctuations about the equilibrium bond length. We added a repulsive potential (the third term in Eq. S1) to avoid unphysical overlaps between the beads.

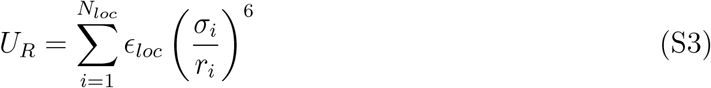

In Eq. S3, *σ*_*i*_ is the sum of the radii of the pair of beads that are interacting. When two backbone beads are interacting then *σ*_*i*_ = 2*σ*_*BB*_, where *σ*_*BB*_ is the radius (=1.9 Å) of the *C*_*α*_ atom. For the interaction between the backbone and sidechain 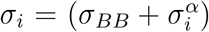. Here, *i* is the residue number and *α* is the type of residue(see Table S1). For sidechain-sidechain interaction 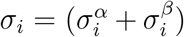 where *α* and *β* are the residue types. The In Eq. S3 we take into account nearest neighbors of a given residue that are separated by less than or equal to two backbone beads, which are not connected by a covalent bond. For example, the backbone of residue 1 would have repulsive interaction with sidechain of residue 2, and both backbone and sidechain beads of the third residue. Similarly, a repulsive interaction exists between the side chain of residue 1 and the backbone and the side chain bead of residues 2 and 3.

Electrostatic interaction are modeled through the Debye-Huckel potential:

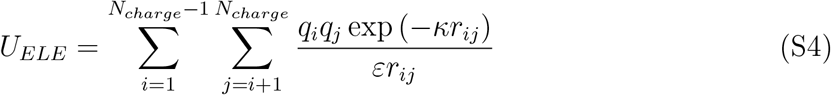

where *q*_*i*_ and *q*_*j*_ are the point charges located at the center of the the side chain beads of charged residues. In the SOP-IDP model, *q*_*i*_ = +1 (for *q*_*i*_ = −1) positively (negatively) charged residues. The inverse Debye length corresponds to the 100 mM NaCl salt concentration. The dielectric constant was taken to be 78.

The backbone-backbone (BB), backbone-side-chain (BS), and side-chain-side-chain (SS) interactions are modeled using the Lennard-Jones (LJ) potentials:

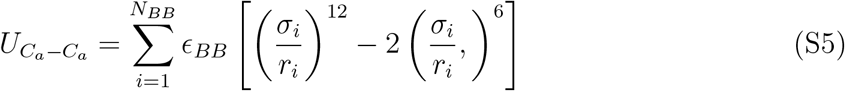

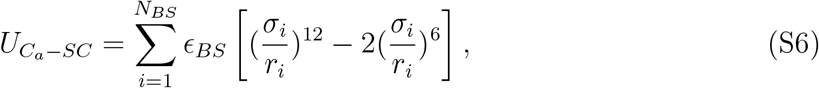

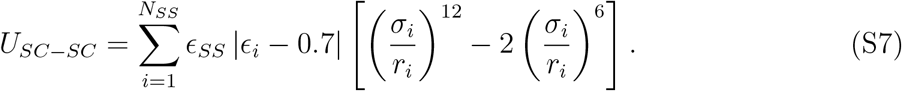

The optimum values of *ϵ*_*BB*_, *ϵ*_*BS*_, and *ϵ*_*BS*_ were determined using a learning procedure.^S1^ The pairwise interactions between the sidechain beads (final term in Eq. S1) account for the sequence-specificity encoded in the SOP-IDP model. The parameter *ϵ*_*i*_ is based on the Betancourt-Thirumalai matrix, ^S3^ which sets the interaction scale between different amino acids.

#### Simulations

To probe the equilibrium ensembles of the monomeric peptides, we carried out underdamped Langevin dynamics simulations using the LAMMPS package.^S4^ The equation of motion for bead *i* is given by:

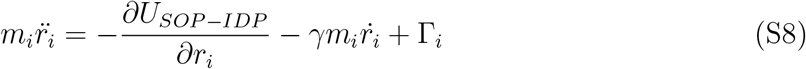

where *m*_*i*_ is the mass of each bead, *r*_*i*_ is the coordinate of the *i*^*th*^ bead, *γ* is the friction coefficient, and Γ_*i*_ is a Gaussian random force which satisfies the fluctuation-dissipation relation, ⟨Γ_*i*_(*t*)Γ_*j*_(*t*′)⟩ = 6k_*B*_T*m*_*i*_*γ*_*i*_*δ*_*ij*_*δ*(*t* − *t*′). The equations of motion were integrated using the velocity-Verlet algorithm, as implemented within the LAMMPS package, using a time-step of 10 fs. All the simulations were carried out at a temperature of 298 K. The friction coefficient *γ* is 0.1 *ps*^−1^. The mass of each bead is 55 g/mol. All the monomers were equilibrated for the first 2 × 10^8^ steps, followed by a production run of at least 8 × 10^8^ steps. To obtain meaningful statistics for the thermodynamic quantities, ten trajectories were generated from different initial condition and we collect at least two million trajectory frames for FUS-LC, FUS-LC-N, FUS-LC-N2, FUS-LC-C and P-FUS (in) systems and 200,000 trajectory frames for P-FUS (out). Because our focus is on equilibrium properties the precise value of *γ* is not relevant.

#### Model for phosphorylation

We mutated the residues at the site of phosphorylation to glutamic acids. Thus, the CG model phosphorylation introduces a negative charge on the side chain. For phosphorylation inside the core-1, we replace the five serine residues S42, S54, S61, S84, S87, and a Threonine T68 with glutamic acid. To model phosphorylation outside the core-1 region, we perform phosphomimetic mutation by mutating T11, T19, and T30 and S112, S117, and S131 outside core-1 with glutamic acid. The interactions between the charged residues as a result of phosphorylation are modeled using Eq. S4.

**Table S1:**
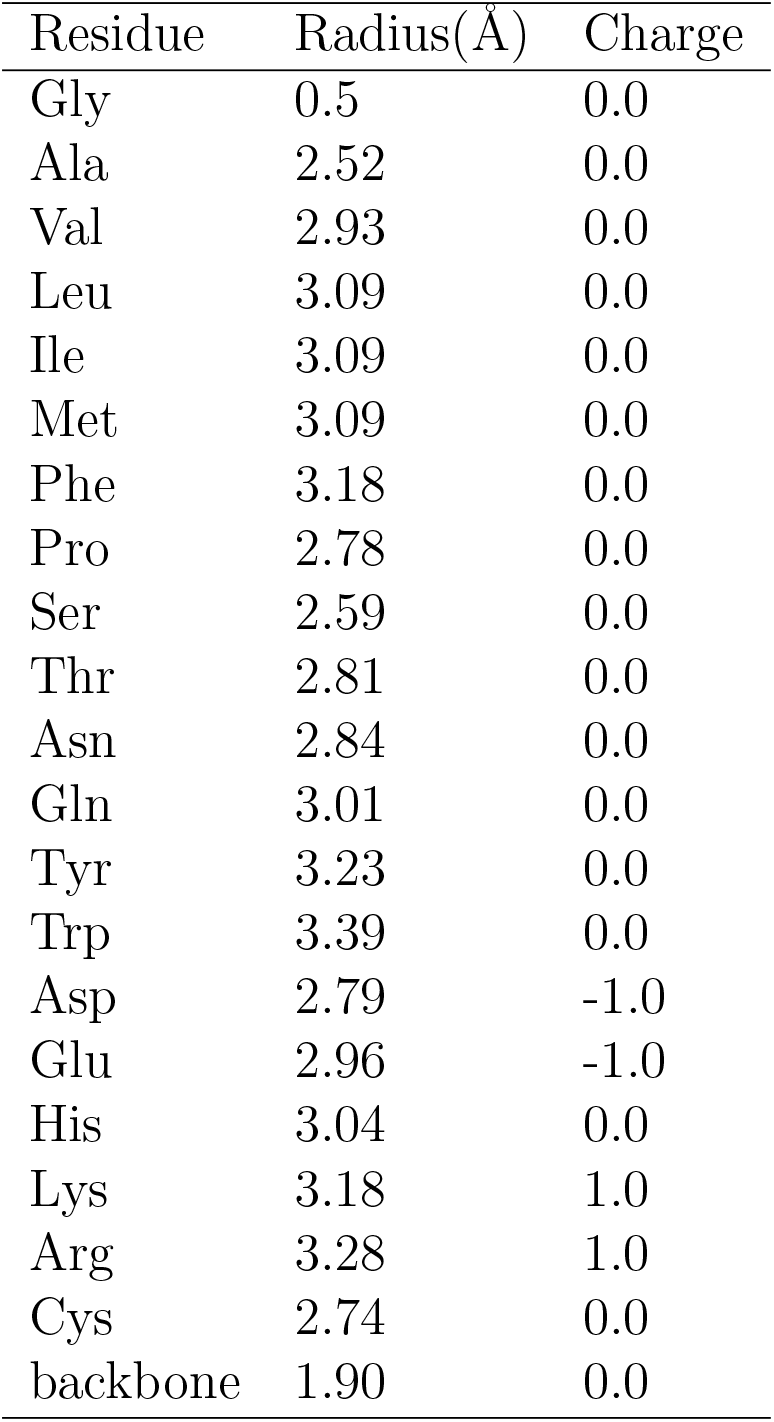
backbone and side chain radius used in the SOP-IDP model

**Table S2:** Energy function parameters for the SOP-IDP model

| Parameters | Value |
| --- | --- |
| k | 20 kcal/mol/$\text{\AA}^2$ |
| $R_o$ | 2 $\text{\AA}$ |
| $\epsilon_{loc}$ | 1 kcal/mol |
| $\epsilon_{BB}$ | 0.12 kcal/mol |
| $\epsilon_{BS}$ | 0.24 kcal/mol |
| $\epsilon_{SS}$ | 0.18 kcal/mol |

### List of FUS-LC mutants and truncated variants used in this study and their FASTA sequences

#### FUS-LC

MASNDYTQQATQSYGAYPTQPGQGYSQQSSQPYGQQSYSGYSQSTDTSG

YGQSSYSSYGQSQNTGYGTQSTPQGYGSTGGYGSSQSSQSSYGQQSSYPG

YGQQPAPSSTSGSYGSSSQSSSYGQPQSGSYSQQPSYGGQQQSYGQQQSYN

PPQGYGQQNQYNSSSGGGGGGGGGGNYGQDQSSMSSGGGSGGGYGNQD

QSGGGGSGGYGQQDRG

#### FUS-LC-N (1-163)

MASNDYTQQATQSYGAYPTQPGQGYSQQSSQPYGQQSYSGYSQSTD

TSGYGQSSYSSYGQSQNTGYGTQSTPQGYGSTGGYGSSQSSQSSYGQ

QSSYPGYGQQPAPSSTSGSYGSSSQSSSYGQPQSGSYSQQPSYGGQQQ

SYGQQQSYNPPQGYGQQNQYNS

#### FUS-LC-N2 (1-108)

MASNDYTQQATQSYGAYPTQPGQGYSQQSSQPYGQQSYSGYSQSTDT

SGYGQSSYSSYGQSQNTGYGTQSTPQGYGSTGGYGSSQSSQSSYGQQ

SSYPGYGQQPAPSS

#### FUS-LC-C (111-214)

GSYGSSSQSSSYGQPQSGSYSQQPSYGGQQQSYGQQQSYNPPQGYG

QQNQYNSSSGGGGGGGGGGNYGQDQSSMSSGGGSGGGYGNQDQS

GGGGSGGYGQQDRG

#### P-FUS (inside core-1)

MASNDYTQQATQSYGAYPTQPGQGYSQQSSQPYGQQSYSGYEQSTD

TSGYGQSEYSSYGQEQNTGYGEQSTPQGYGSTGGYGSEQSEQSSYG

QQSSYPGYGQQPAPSSTSGSYGSSSQSSSYGQPQSGSYSQQPSYGGQ

QQSYGQQQSYNPPQGYGQQNQYNSSSGGGGGGGGGGNYGQDQS

SMSSGGGSGGGYGNQDQSGGGGSGGYGQQDRG

#### P-FUS (outside core-1)

MASNDYTQQAEQSYGAYPEQPGQGYSQQSEQPYGQQSYSGYSQSTD

TSGYGQSSYSSYGQSQNTGYGTQSTPQGYGSTGGYGSSQSSQSSYG

QQSSYPGYGQQPAPSSTSGEYGSSEQSSSYGQPQSGSYEQQPSYGG

QQQSYGQQQSYNPPQGYGQQNQYNSSSGGGGGGGGGGNYGQDQS

SMSSGGGSGGGYGNQDQSGGGGSGGYGQQDRG

### Analyses

#### Structural overlap order parameter

We define a structural overlap parameter, ^S5^ *χ*, as a measure of the similarity of the monomer conformation to the one in the fibril. We also calculated pair distances within a monomer in the simulated conformations and compared them with those in the fibril state. If the pair distances are within 2 Å of each other, we consider the pair distance to be similar. The fraction of the number of such pairs to the total number of pairs is the value of the order parameter. The structural overlap has a bound 0 *≤ χ ≤* 1. We calculated *χ*_1_ for core-1 (39-95 residues) whereas *χ*_2_ is computed for the 112-150 region. We defined *χ* as,

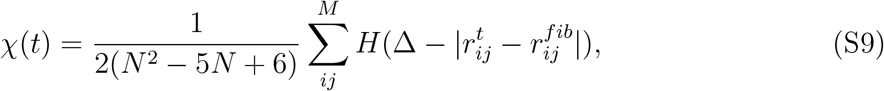

where *N* is the number of residues in the core of the fibril, *H*(*x*) is the Heaviside function, and 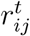 and 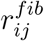 are the pair distances for the simulated conformations and the experimental fibril structure, respectively. We set Δ = 2 Å. The total number of pair distances is M = 2(*N*^2^ − 5*N* + 6) as shown in the caption of Fig. S1. This factor ensures that in the fibril state *χ* = 1.

#### Threshold value of *χ*_*c*_ to identify core-1 and core-2 fibril like conformations

We identified the N* states using *χ*_1_ (core-1) and *χ*_2_ (core 2). The distances between the fibril monomers are for the core-1 (residues 39-95) were calculated using the experimental structure having the S-bend topology (PDB ID: 5W3N). For core-2 (residues 112-150) we used the structure with the U-bend topology (PDB ID: 6XFM). We used *χ*_*c*_ *>*0.215 as representing the N* state for both core-1 and core-2 fibrils. We arrived at the *χ*_*c*_ using the following method. For the core-1 region, we identified the key contacts among amino acid residues that maintain the S-shape of the FUS fibril. We restrained the distances between these residues to the values in the fibril monomer using a harmonic bond between backbone residues. The strength of the harmonic bond is kept at 50 kcal/mol Å^2^. The five harmonic constraints are between backbone residues 76-83, 69-92, 44-78, 55-62, 47-70. The corresponding distances are 7.091, 6.15, 6.68, 6.99 and 6.81 Å. From 2 *×* 10^6^ conformations obtained from the constrained simulations, we calculated the distribution of the *χ*. We find that the peak for the constrained core-1 conformations is *χ* = 0.215, which was used as the threshold value both the cores of the fibrils.

**Figure S1:**
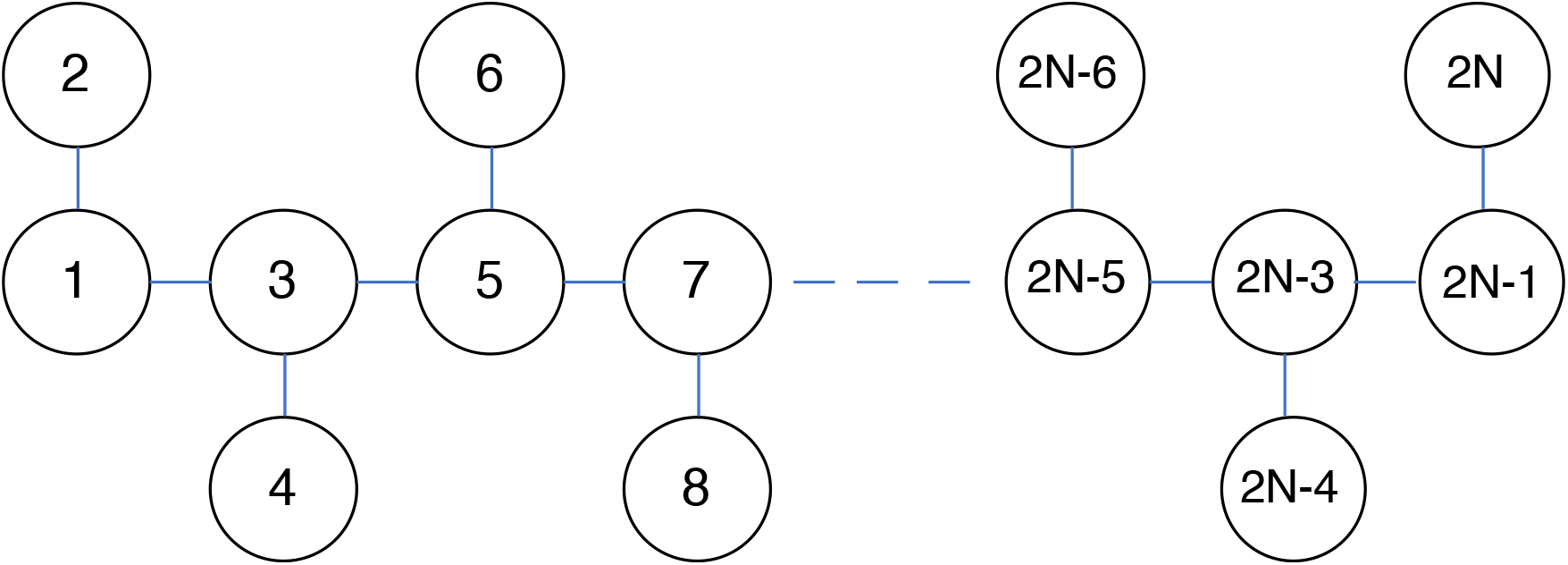
List of pair distances for the structural overlap parameter. There are 2N beads in the simulation for the protein-containing N residue. The total number of possible pair distances is 2N(2N-1)/2. However, we excluded covalent bonds and neighbors belonging to two nearest residues. So the pair distances that are considered for bead 1 are (1,7),(1,8),(1,9)….(1,2N). The pair distances for bead 2 are (2,7),(2,8),(2,9). (2,2N). For bead 3, pair distances are (3,9),(3,10),(3,11)….(3,2N), for bead 2N-8, pair distances are (2N-8,2N-1),(2N-8,2N), for bead 2N-7, pair distances are (2N-7,2N-1), (2N-7,2N). The total number of pairs for calculating the structure overlap parameter is 2(*N*^2^ − 5*N* + 6).

#### Contact map

If the distance between side chains of two residues is less than or equal to 10 Å, the residues are assumed to be in contact. Rather than using a step function, we assessed if a contact is formed using the logistic function:

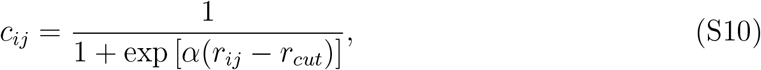

where *r*_*cut*_ = 10 Å, and *α* = 50 Å^−1^. Here, *r*_*ij*_ is the distance between two non-bonded beads. The large *α* value ensures that *c*_*ij*_ changes abruptly from zero to unity over a narrow range of |*r*_*ij*_ − *r*_*cut*_|. We calculated the average contact map using the 8195 (666) N^∗^ conformations that belong to core-1 (core-2).

#### Relative stability of core-1 vs core-2

We counted the conformation that has a more significant structural overlap with core-1 and core-2 using *χ*. Using the number of conformations that are similar to core-1 and core-2, we computed the average potential energy *U*_*SOP* −*IDP*_ for the 8195 core-1 and 666 core-2 conformations obtained from the full-length FUS-LC simulation. We use the block average method to compute the error bars. The average total energy of core-1 and core-2 is 200.03 ± 0.11 and 202.58 ± 0.36 kcal/mol, respectively. The difference in the total energy between core-1 and core-2 energy is −2.55 ± 0.38 kcal/mol. The free energy difference between core-1 and core-2 is Δ*G* = *G*_1_ − *G*_2_ = −*k*_*B*_*T ×* ln(*P*_*core*−1_*/P*_*core*−2_) = −0.593 *×* ln(8165 ± 103*/*666 ± 29) = −1.47 ± 0.005 kcal/mol. The *T* Δ*S* ≈ −1.08 ± 0.38 kcal/mol and the entropy difference Δ*S* is *S*_1_ − *S*_2_ = −3.62 ± 1.27 cal/mol/K. Thus, the simulations predict that core-1 is free energetically preferred over core-2. We should note that computation of *Z*-scores associated with the energies also clearly show that core-1 has lower energy than core-2.

#### All-atom reconstruction of coarse-grained structures

The all-atom reconstruction was carried out using the Protein Chain Reconstruction Algorithm (PULCHRA). ^S6^ The reconstructed structure was further optimized using the AMBER ff12SB force field, in conjunction with the Generalized Born solvent model (GB-OBC), ^S7^ as implemented within the AMBER12 code.^S8^ During geometry optimization, the C_*α*_ atoms were restrained to prevent significant deviations from the topology of the coarse-grained structure.

**Figure S2:**
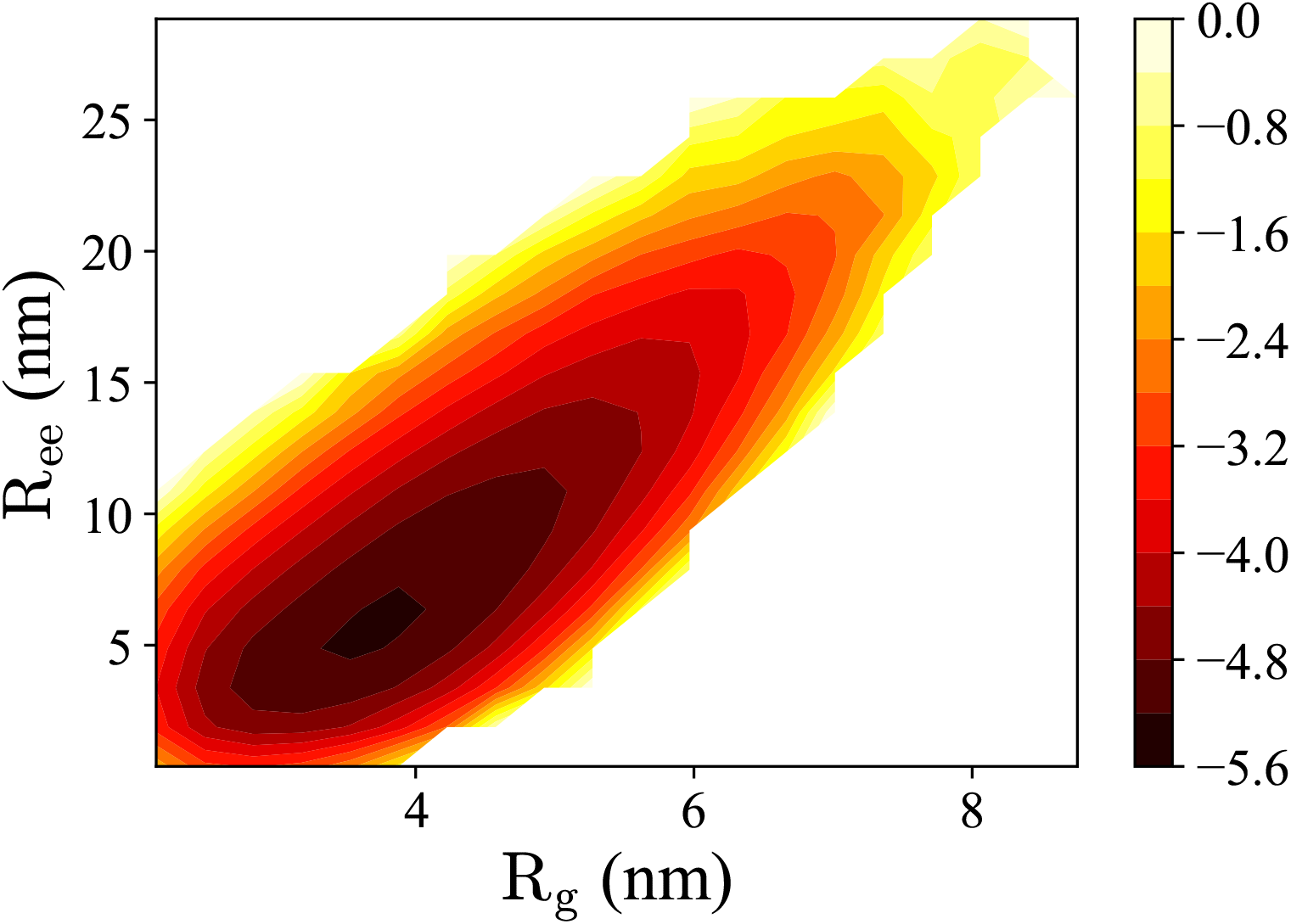
Free energy landscape of the FUS-LC monomer projected onto the space of radius of gyration (*R*_*g*_) and end-to-end distance (*R*_*ee*_). The landscape is essentially featureless, and lacks any discernible metastable states.

**Figure S3:**
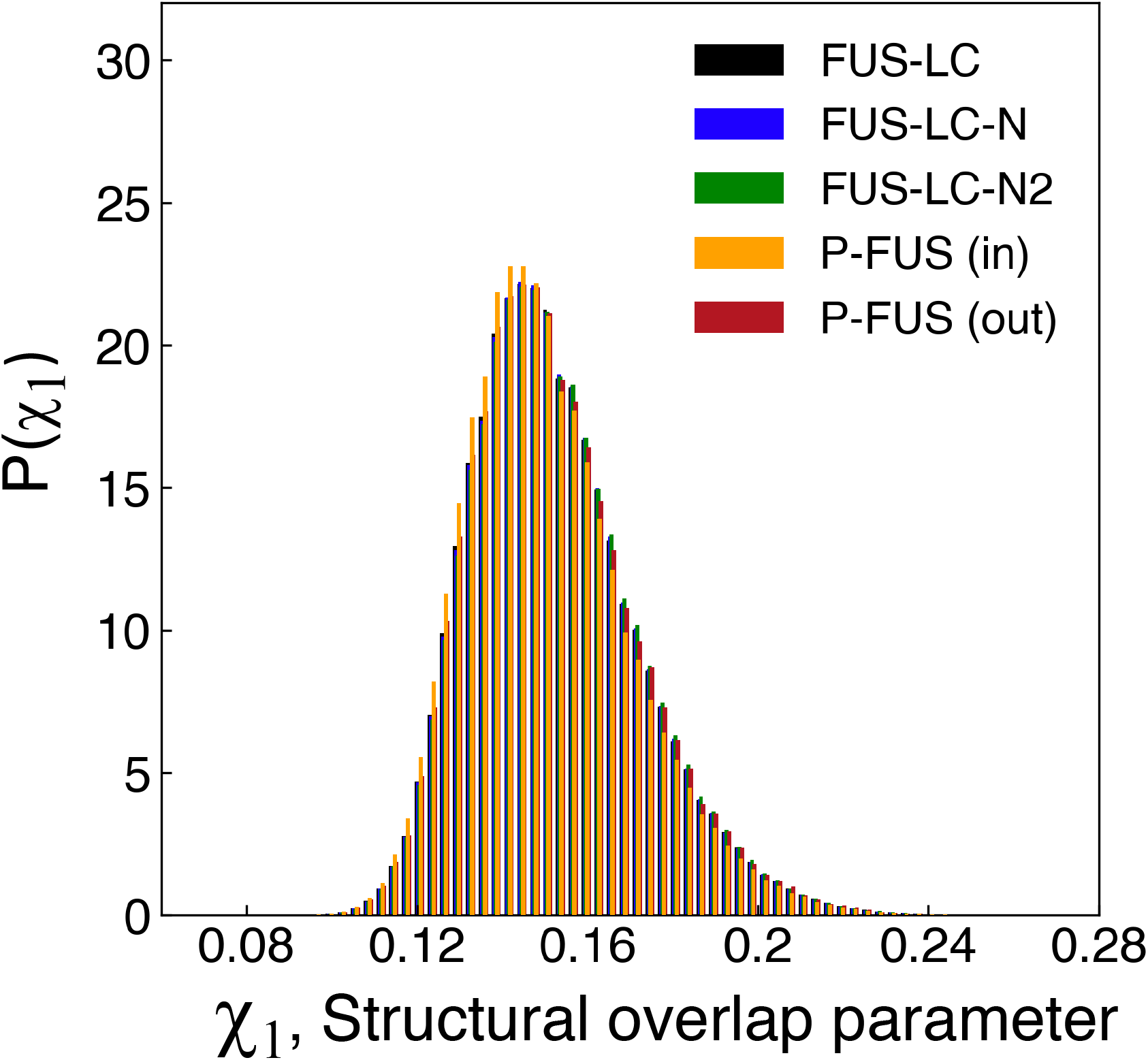
Distribution of the structural overlap parameter (*χ*_1_) for FU-LC (1-214) (black), FUS-LC-N (1-163) (blue), FUS-LC-N2 (green), P-FUS (in) (phosphorylation inside the core-1) (Orange), P-FUS (out) (phosphorylation outside the core-1 region) (maroon). We shifted the x-axis such that distance between two bars in the distribution is one-tenth of the bin width to make the distributions visible.

**Figure S4:**
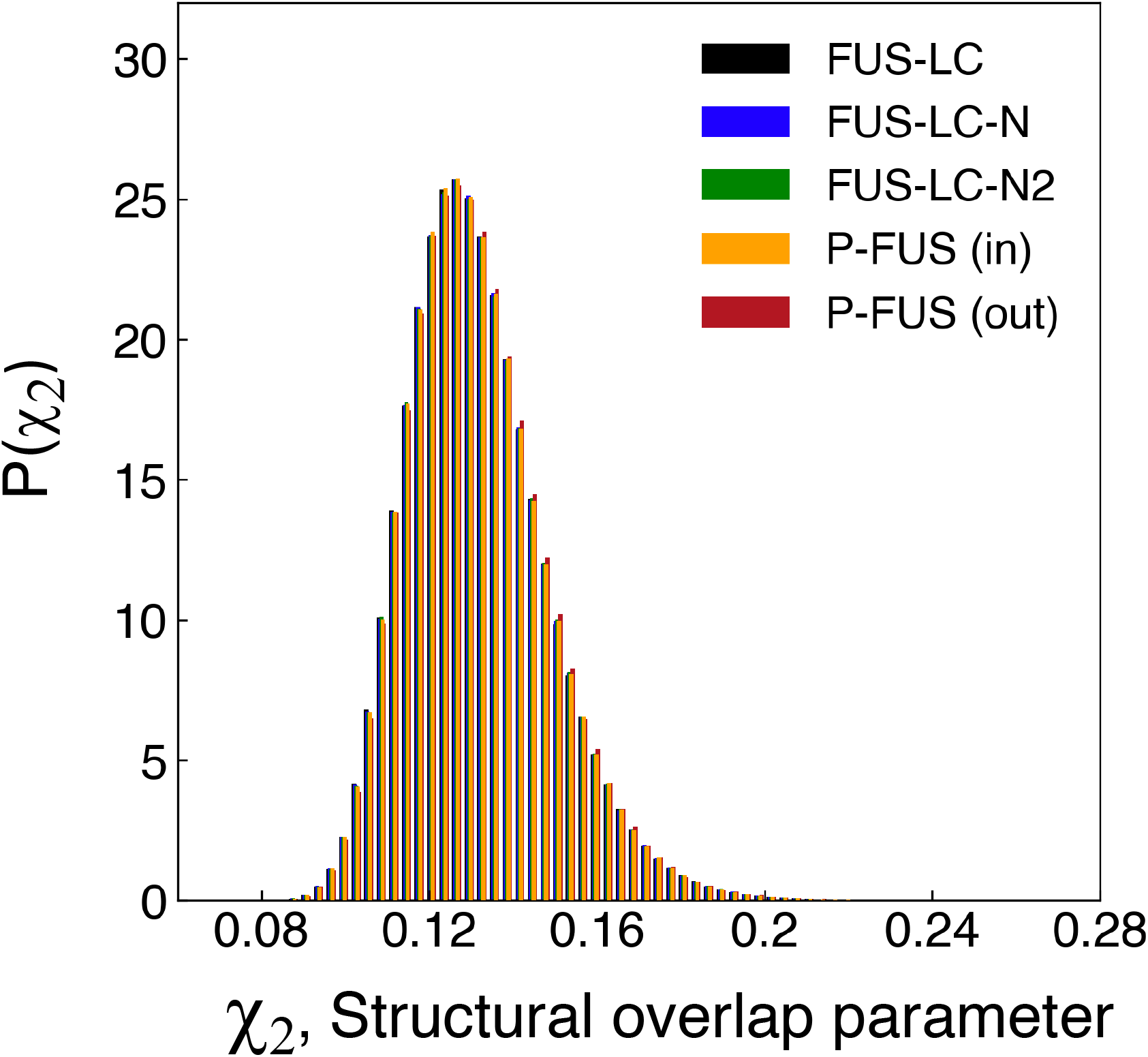
Distribution of the structural overlap parameter (*χ*_2_) for FU-LC (1-214) (black), FUS -LC-N (1-163) (blue), FUS-LC-N2 (green), P-FUS (in) (phosphorylation inside the core-1) (Orange), P-FUS (out) (phosphorylation outside the core-1 region) (maroon). We shifted the x-axis such that distance between two bars of the distribution is one-tenth of the bin width to make the distribution visible.

**Figure S5:**
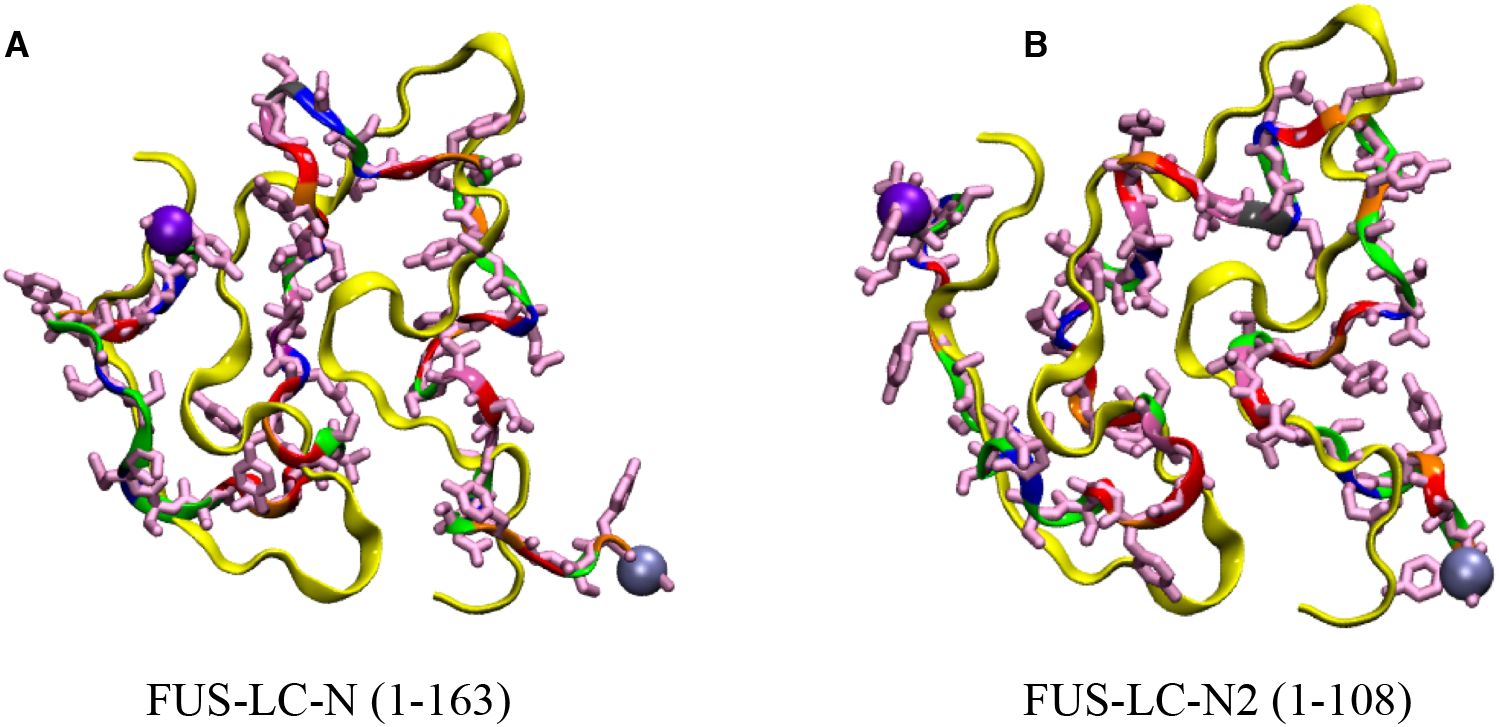
Representative N^∗^ states from the conformational ensembles of the FUS-LC-N (A), and the FUS-LC-N2 (B) sequences, having the core-1 topology. In both cases, the S-bend morphology is largely preserved, suggesting that both sequences could form the core-1 fibril.

**Figure S6:**
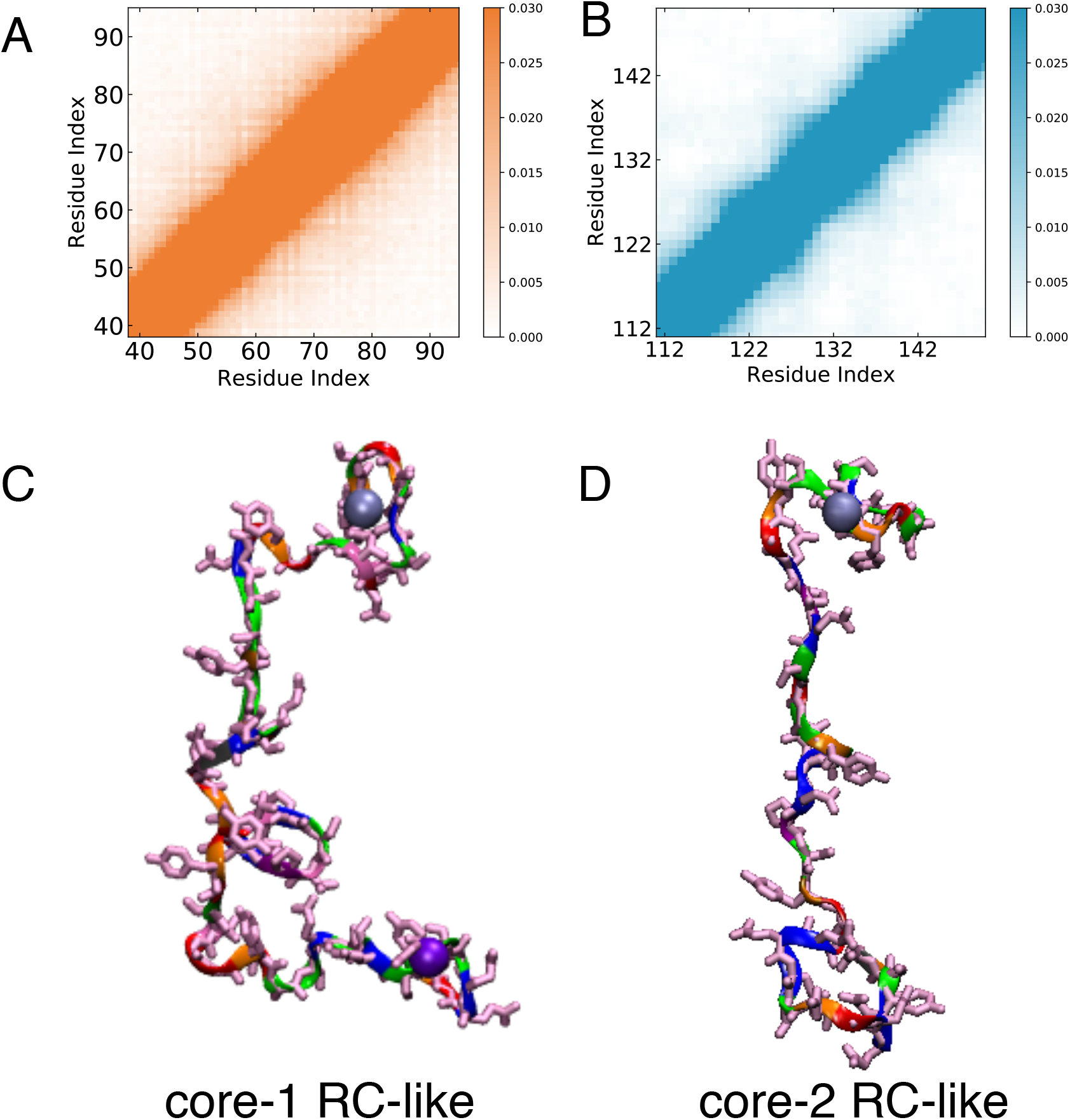
Contact maps and structures of core-1 and core-2 region from conformations that are not the N* state A) and B) shows the contact map of FUS-LC in the core-1 and core-2 region. We compute the contact map for all the conformations that do not contain the N^∗^ state corresponding to core-1 and core-2 respectively C) and D) shows a snapshot of core-1 and core-2 region that corresponds to random-coil (RC) like conformation.

